# Chromosome-level assembly of the *Caenorhabditis remanei* genome reveals conserved patterns of nematode genome organization

**DOI:** 10.1101/2019.12.31.892059

**Authors:** Anastasia A. Teterina, John H. Willis, Patrick C. Phillips

## Abstract

The nematode *Caenorhabditis elegans* is one of the key model systems in biology, including possessing the first fully assembled animal genome. Whereas *C. elegans* is a self-reproducing hermaphrodite with fairly limited within-population variation, its relative *C. remanei* is an outcrossing species with much more extensive genetic variation, making it an ideal parallel model system for evolutionary genetic investigations. Here, we greatly improve on previous assemblies by generating a chromosome-level assembly of the entire *C. remanei* genome (124.8 Mb of total size) using long-read sequencing and chromatin conformation capture data. Like other fully assembled genomes in the genus, we find that the *C. remanei* genome displays a high degree of synteny with *C. elegans* despite multiple within-chromosome rearrangements. Both genomes have high gene density in central regions of chromosomes relative to chromosome ends and the opposite pattern for the accumulation of repetitive elements. *C. elegans* and *C. remanei* also show similar patterns of inter-chromosome interactions, with the central regions of chromosomes appearing to interact with one another more than the distal ends. The new *C. remanei* genome presented here greatly augments the use of the *Caenorhabditis* as a platform for comparative genomics and serves as a basis for molecular population genetics within this highly diverse species.

## Introduction

The free-living nematode *Caenorhabditis elegans* is one of the most-used and best-studied model organisms in genetics, developmental biology, and neurobiology (Brenner 1973; Brenner 1974; Blaxter 1998). *C. elegans* was the first multicellular organism with a complete genome sequence (*C. elegans* Sequencing Consortium 1998), and the *C. elegans* genome currently has one of the best described functional annotations among metazoans, as well as possessing hundreds of large-scale datasets focused on functional genomics (Gerstein *et al*. 2010). The genome of *C. elegans* is compact, roughly one hundred megabases (100.4 Mb is the “classic” N2 assembly (*C. elegans* Sequencing Consortium 1998); 102 Mb is the V2010 strain genome (Yoshimura *et al*. 2019)), and consists of six holocentric chromosomes, five of which are autosomes and one that is a sex chromosome (X). All chromosomes of *C. elegans* have a similar pattern of organization: a central region occupying about half of the chromosome that has a low recombination rate, low transposon density, and high gene density, and the “arms” display the characteristics exactly opposite to this (Waterston *et al*. 1992; Barnes *et al*. 1995; Rockman and Kruglyak 2009).

About 65 species of the *Caenorhabditis* genus are currently known (Kiontke *et al*. 2011), and for many of them genomic sequences are available (Stevens *et al*. 2019) http://www.wormbase.org/ and https://evolution.wormbase.org). Most of the *Caenorhabditis* nematodes are outcrossing species with females and males (gonochoristic), but three species, *C. elegans, C. briggsae*, and *C. tropicalis*, reproduce primarily via self-fertilizing (“selfing”) hermaphrodites with rare males (androdioecy) (Kiontke *et al*. 2011). *Caenorhabditis* species have the XX/XO sex determination: females and hermaphrodites carry two copies of the X chromosomes while males have only one X chromosome (Pires-daSilva 2007).

*Caenorhabditis remanei* is an obligate outcrossing nematode, a member of the ‘Elegans’ supergroup, which has become an important model for natural variation (Jovelin *et al*. 2003; Reynolds and Phillips 2013), experimental evolution (Sikkink *et al*. 2014a; Castillo *et al*. 2015; Sikkink *et al*. 2015; Sikkink *et al*. 2019) and population genetics (Graustein *et al*. 2002; Cutter *et al*. 2006; Cutter and Charlesworth 2006; Dolgin *et al*. 2007; Jovelin and Phillips 2009; Dey *et al*. 2012). Whole-genome data is available for three strains of *C. remanei* (Table 1), but all of these assemblies are fragmented. To improve a resolution of experimental studies and analyze chromosome-wide patterns of genome organization, recombination and diversity, the complete assembly for this species is required. We generated a chromosome-level genome assembly of the *C. remanei* PX506 inbred strain using a long-read/Hi-C approach and used this new chromosome-level resolution in a comparative framework to reveal global similarities in genome organization and spatial chromosome interactions between *C. elegans* and *C. remanei*.

## Materials and Methods

### Nematode strains

Nematodes were maintained under standard lab conditions as described (Brenner 1974). *C. remanei* isolates were originally derived from individuals living in association with terrestrial isopods (Family *Oniscidea)* collected from Koffler Scientific Reserve at Jokers Hill, King City, Toronto, Ontario as described in (Sikkink *et al*. 2014b). Strain PX393 was founded from a cross between single female and male *C. remanei* individuals isolated from isopod Q12. This strain was propagated for 2-3 generations before freezing. PX506, the source of the genome described here, is an inbred strain derived from PX393 following sib-mating for 30 generations to reduce within-strain heterozygosity. This strain was subsequently frozen and subsequently recovered at large population size for several generations before any subsequent experimental analysis.

### Sequencing and genome assembly of the *C. remanei* reference

Strain PX506 was grown on twenty 110mm plates until its entire *E. coli* food source (strain OP50) was consumed. Worms were washed 5X in M9 using 15 ml conical tubes and spun at low RPM to concentrate. The worm pellet was flash frozen and genomic DNA was isolated using the Genomic-tip 100/G column (Qiagen). Four micro-grams (average size 23Kb) was frozen and shipped to Dovetail genomics (Santa Cruz, CA, USA, https://dovetailgenomics.com) along with frozen whole animals for subsequent PacBio and Hi-C analysis. The *C. remanei* PX506 inbred strain was sequenced and assembled by Dovetail Genomics. The primary contigs were generated from two PacBio SMRT Cells using the FALCON assembly (Chin *et al*. 2016) followed by the Arrow polishing (https://github.com/PacificBiosciences/GenomicConsensus). The final scaffolds were constructed with Dovetail Hi-C library sequences and the HiRise software pipeline (Putnam *et al*. 2016). Additionally, we performed whole-genome sequencing of PX506 strain using the Nextera kit (Illumina) for 100bp paired-end read sequencing on the Illumina Hi-Seq 4000 platform (University of Oregon Sequencing Facility, Eugene, OR).

We then performed a BLAST search (Altschul *et al*. 1990) against the NCBI GenBank nucleotide database (Benson *et al*. 2012) and filtered any scaffolds (e-value < 1e-15) of bacterial origin. Short scaffolds with good matches to *Caenorhabditis* nematodes were aligned to 6 chromosome-sized scaffolds by GMAP v.2018-03-25 (Wu and Watanabe 2005) and visualized in IGV v.2.4.10 (ThorvaldsdÓttir *et al*. 2013) to examine whether they represent alternative haplotypes.

The final filtered assembly was compared to the “recompiled” version of *C. elegans* reference genome, generated from strain VC2010, a modern strain derived from the classical N2 strain (Yoshimura *et al*. 2019) and *C. briggsae* genomes (available under accession numbers PRJEB28388 from the NCBI Genome database and PRJNA10731 form WormBase WS260) by MUMmer3.0 (Kurtz et al. 2004). The names and orientations of the *C. remanei* chromosomes were defined by the longest total nucleotide match in proper orientation to *C. elegans* chromosomes. Dot plots with these alignments were plotted using the ggplot2 package (Wickham 2016) in R (R Core Team 2018). The completeness of the *C. remanei* genome assembly was assessed by BUSCO v.3.0.2 (Simão *et al*. 2015) with Metazoa odb9 and Nematoda odb9 databases. Results were visualized with generate_plot_xd_v2.py script (https://github.com/xieduo7/my_script/blob/master/busco_plot/generate_plot_xd_v2.py). The mitochondrial genome was generated using a reference mitochondrial genome of *C. remanei* (KR709159.1) from the NCBI database (http://www.ncbi.nlm.nih.gov/nucleotide/) and Illumina reads of the *C. remanei* PX506 inbred strain. The reads were aligned with bwa mem v.0.7.17 (Li and Durbin 2009); filtered with samtools v.1.5 (Li *et al*. 2009a). We marked PCR duplicates in the mitochondrial assembly with MarkDuplicates from picard-tools v.2.0.1 (http://broadinstitute.github.io/picard/); realigned indels and called variants with IndelRealignment and HaplotypeCaller in the haploid mode from GATK tools v.3.7 (McKenna *et al*. 2010), filtered low-quality sites, and then used bcftools consensus v.1.5 (Li 2011) to generate the new reference mitochondrial genome. To estimate the residual heterozygosity throughout the rest of the genome, we implemented a similar readmapping protocol but used the default parameters to call genotypes and then filtered variants using standard hard filters.

### Repeat masking in *C. remanei* and *C. elegans*

For the repeat masking, we created a comprehensive repeat library (Coghlan *et al*. 2018)’ see also instructions at http://avrilomics.blogspot.com) and masked sequencespecific repeat motifs, as described in (Woodruff and Teterina 2019). *De novo* repeat discovery was performed by RepeatModeler v.1.0.11, (Smit and Hubley 2008) with the ncbi engine. Transposon elements were detected by transposonPSI (http://transposonpsi.sourceforge.net), with sequences shorter than 50 bases filtered out. Inverted transposon elements were located with detectMITE v.2017-04-25 (Ye *et al*. 2016) with default parameters. tRNAs were identified with tRNAscan-SE v.1.3.1 (Lowe and Eddy 1997) and their sequences were extracted from a reference genome by the getfasta tool from the BEDTools package v.2.25.0 (Quinlan and Hall 2010). We searched for LTR retrotransposons as described at http://avrilomics.blogspot.com/2015/09/ltrharvest.html, by LTRharvest and LTRdigest from GenomeTools v.1.5.11 (Gremme *et al*. 2013) with domains from the Gypsy Database (Llorens *et al*. 2010) and several models of Pfam protein domains (Finn *et al*. 2015) listed in Supplementary Tables B1 and B2 of (Steinbiss *et al*. 2009); to filter LTRs we used two scripts, https://github.com/satta/ltrsift/blob/master/filters/filter_protein_match.lua and get_ltr_retrotransposon_seqs.pl from https://gist.github.com/avrilcoghlan.

Additionally, we uploaded nematode repeats from the Dfam database (Hubley *et al*. 2015) using the *queryRepeatDatabase.pl* script from the RepeatMasker v.4.0.7 (Smit *et al*. 2015) utils with “-species rhabditida” option, and *C. elegans* and ancestral repetitive sequences from Repbase v.23.03, (Bao *et al*. 2015). We then combined all repetitive sequences obtained from these tools and databases to one redundant repeat library. We clustered those sequences with less than 80% identity by uclust from the USEARCH package v.8.0, (Edgar 2010) and classified them by the RepeatMasker Classify tool v.4.0.7, (Smit *et al*. 2015). Potential protein match with *C. remanei* (PRJNA248911) or *C. elegans* protein sequences (PRJNA13758) from WormBase W260 were detected with BLASTX (Altschul *et al*. 1990). The repetitive sequences classified as “unknown” and having blast hits with E-value≤0.001 with known protein-coding genes were removed from the final repeat libraries.

For *C. remanei*, the final repeat library was used by RepeatMasker with “-s” and “-gff’ options. Additional round of masking was performed with “-species caenorhabditis” option. The genome was also masked with the redundant repeat library acquired before the clustering step. Regions that were masked with redundant library but not masked with the “final” library were extracted using BEDTools subtract and classified by RepeatMasker Classify. Additionally, we checked the depth coverage with the Illumina reads in these regions, as regions classified as a known type of repeat and displaying coverage more than 70 were masked in the reference genome by BEDTools maskfasta. The masked regions were extracted to a bed file with a bash script (https://gist.github.com/danielecook/cfaa5c359d99bcad3200), and the same regions were soft masked by BEDTools maskfasta with “-soft” option.

Using the same approach, we masked the *C. elegans* reference of N2 strain (PRJNA13758 from WormBase W260) and then extracted all regions that were masked in the “official” masked version of the genome but not masked by our final repeat library. These regions were extracted, classified by RepeatMasker with default parameters, and searched against *C. elegans* proteins with BALSTX algorithm and the *C. elegans* reference genome with BLASTN. Regions with “unknown” class and a match with *C. elegans* proteins (see above) were removed. Regions with more than 5 matches and evalue ≤0.001 with the *C. elegans* genome were added to the final database and used to mask the *C. elegans* reference genome generated from strain VC2010 (Yoshimura *et al*. 2019). The same regions were soft-masked with BEDTools maskfasta.

### Full-length transcript sequencing

We used single molecule long-read RNA sequencing (Iso-Seq) to obtain high-quality transcriptomic data. We used the Clonetech SMARTer PCR cDNA Synthesis kit for cDNA synthesis and PCR amplification with no size selection starting with 500ng of total RNA from a mixed staged population of *C. remanei* strain PX506 (Clonetech, Cat#634925). PacBio library generation was performed on-site at the University of Oregon genomics facility and sequenced on a PacBio Sequel I platform utilizing four SMRT cells of data.

We generated circular consensus reads by ccs tool with “--noPolish --minPasses 1” options from PacBio SMRT link tools v.5.1.0 (https://www.pacb.com/support/software-downloads/) and obtained full-length transcripts with lima from the same package with “--isoseq --no-pbi” options. Then trimmed reads from all SMRT cells were merged together, clustered and polished with isoseq3 tools v.3.2 (https://github.com/PacificBiosciences/IsoSeq), and mapped to the *C. remanei* reference genome with GMAP (Wu and Watanabe 2005). Redundant isoforms were collapsed by collapse_isoforms_by_sam.py from Cupcake ToFU (https://github.com/Magdoll/cDNA_Cupcake). The longest ORF were predicted with TransDecoder v.5.0.1 (Haas *et al*. 2013) and used as CDS hints in the genome annotation (see below).

### Genome annotation

We performed de novo annotation of the *C. remanei* genome using the following hybrid approach. For *ab initio* gene prediction, we applied the GeneMark-ES algorithm v.4.33 (Ter-Hovhannisyan *et al*. 2008) with default parameters. De novo gene prediction with the MAKER pipeline v.2.31.9 (Holt and Yandell 2011) was carried out with *C. elegans* (PRJNA13758), *C. briggsae* (PRJNA10731), *C. latens* (PRJNA248912) proteins from Wormbase 260, excluding the repetitive regions identified above. To implement gene prediction using the BRAKER pipeline v.2.1.0 (https://github.com/Gaius-Augustus/BRAKER), we included RNA-seq from our previous *C. remanei* studies (SRX3014311 and SRP049403).

Annotations from BRAKER2, MAKER2, and GeneMark-ES were combined in EVidenceModeler v.1.1.1 (Haas *et al*. 2008) with weights 6,3, and 1, correspondingly. CDS from the EVidenceModeler results were used to train AUGUSTUS version 3.3 (Stanke *et al*. 2006) as described on http://bioinf.uni-greifswald.de/augustus/binaries/tutorial/training.html. Then models were optimized and re-trained again. Then we created a file with extrinsic information with factor 1000 and malus 0.7 for CDS, all other options as in “extrinsic.E.cfg” for annotation with est database hits from the AUGUSTUS supplementary files. The final annotation was executed with Iso-seq data as the hints file and EVidenceModeler-trained models with singlestrand=true --gff3=on --UTR=off”. Scanning for known proteins domains and the functional annotation was conducted with InterProScan v.5.27-66.0 (Quevillon *et al*. 2005). We validated/filtered final gene models according to coverage with RNA-seq and Iso-seq data, matches with known *Caenorhabditis* proteins and protein/transposon domains.

We identified 1-to-1 orthologs of *C. remanei* and *C. elegans* proteins using orthofinder2 (Emms and Kelly 2018), for *C. elegans* we used only proteins validated in the VC2010 (Yoshimura *et al*. 2019). The identity of the proteins was estimated by pairwise global alignments using calc_pc_id_between_seqs.pl script (https://gist.github.com/avrilcoghlan/5311008). Gene synteny plots were made in R with a custom script (synteny_plot.R).

### Genome activity and features

We studied patterns of genome activity in *C. elegans* and *C. remanei* using *C. elegans* Hi-C data from the (Brejc *et al*. 2017) study (SRR5341677-SRR5341679) and the Hi-C reads produced in the current study, as well as available RNA-seq data from the L1 larval stage for *C. elegans* (SRR016680, SRR016681, SRR016683) and *C. remanei* (SRP049403). Hi-C reads were mapped to the reference genomes with bwa mem and RNA-seq read with STAR v.2.5 (Dobin *et al*. 2013) using the default parameters and gene annotations; to count reads for transcripts we used htseq-count form the HTSeq package v.0.9.1 (Anders *et al*. 2015) and corrected by the total lengths of the gene CDS (RPKM) using a bash script (https://gist.github.com/darencard/fcb32168c243b92734e85c5f8b59a1c3) and a custom R script (RNA_seq_R_analysis_and_figures.R). For Hi-C interactions, we applied the Arima pipeline (https://github.com/ArimaGenomics/mapping_pipeline), BEDTools, and a custom bash-script (Hi-C_analysis_with _ARIMA.sh and Hi-C_R_analysis_and_figures.R).

We calculated the fraction of exonic/intronic DNA and the number of genes per 100 kb windows from the genome annotations using BEDtools coverage tool. GC-content and the percent of repetitive regions were estimated, correspondingly, from the un-masked and hard masked genomes via BEDtools nuc also on 100 Kb windows by a custom script (get_genomic_fractions.sh). For the formal statistical tests, we defined chromosome “centers” to be the central half of a chromosome and the “arms” to be the peripheral one quarter of each length on either side of the center. To measure the positional effect of these genomic features, we conducted the Cohen’s d effect size test with package “lsr” in R (Navarro 2013) and calculated statistical differences by Wilcoxon-Mann-Whitney test using basic R (see a custom script fractions_stats_and_figures.R).

### Data Availability

Strain PX506 is available from the Caenorhabditis Genetic Center (CGC). All raw sequencing data generated in this study have been submitted to the NCBI BioProject database (https://www.ncbi.nlm.nih.gov/bioproject/) under accession number PRJNA577507. The reference genome is available at the NCBI Genome database (https://www.ncbi.nlm.nih.gov/genome/) under accession number WUAV00000000. Supplementary custom scripts to estimate statistics, and generate main and supplementary figures are available on GitHub (https://github.com/phillips-lab/C.remanei_genome). Supplementary figures, tables, and data are on FigShare (https://figshare.com/projects/Supplementary_Tables_and_Figures_of_Chromosome-level_assembly_of_the_Caenorhabditis_remanei_genome_reveals_conserved_patterns_of_nematode_genome_organization_/73518).

## Results

### New reference genome assembly and annotation

We generated a high-quality chromosome-level assembly of the *C. remanei* PX506 inbred line with deep PacBio whole-genome sequencing (~100x coverage by 1.3M reads) and Hi-C (~900x with 418M of paired-end Illumina reads). Assembly of the PacBio sequences resulted in a 135.85 Mb of genome and bacterial sequences with 298 scaffolds. The HI-C data dramatically improved the PacBio assembly, the HiRise scaffolding increased the N50 from 4.042 Mb to 21.502 Mb by connecting scaffolds from the PacBio assembly together, resulting in 235 scaffolds (see the summary statistics in Table 1 and Supplementary Tables S1, S2, and S3). After the filtering of scaffolds of bacterial origin, 6 chromosome-sized scaffolds were obtained, as expected. Additionally, there were 180 short scaffolds that are alternative haplotypes or unplaced scaffolds (the average length is 31,169 nt with standard deviation of 48,700, and the median length of 19,076 nt). Because only the long-sized fraction of total DNA was selected in the long-read library, the mitochondrial DNA was not covered by PacBio sequencing. The mitochondrial genome was therefore generated independently using the Illumina whole-genome data of the reference strain (see Methods). The total length of the new *C. remanei* reference genome without alternative haplotypes is 124,870,449 bp, which is very close in size to previous assemblies of other *C. remanei* strains (Table 1). After 30 generations of inbreeding, the residual heterozygosity of the PX506 line remained at 0.03% (a 100-fold decrease relative to population-level variability (Dey *et al*. 2012).

**Table 1.** Available genome assemblies of *C. remanei.*

| Strain | NCBI ID | Total<br>size<br>(Mb) | Number<br>of<br>scaffold<br>s | Scaffold<br>N50<br>(Mb) | Scaffold<br>L50 | GC% | Number<br>of<br>genes |
| --- | --- | --- | --- | --- | --- | --- | --- |
| PB4611 | GCA_000149515.1 | 145.44<br>3 | 3670 | 0.435 | 70 | 38.5<br>0 | 32412 |
| PX356 | GCA_001643735.2 | 118.54<br>9 | 1591 | 1.522 | 10 | 35.9<br>0 | 24977 |
| PX439 | GCA_002259225.1 | 124.54<br>2 | 912 | 1.765 | 13 | 35.3<br>0 | 24867 |
| PX506* | XXXXX | 124.87<br>0 | 7 | 21.502 | 3 | 37.9<br>6 | 26308 |

\* - our study.

To assess the quality of the new reference we performed a standard BUSCO analysis (Simão *et al*. 2015). The new assembly of PX506 presented here has 975 of 982 BUSCO genes for completeness 97.9% based on Nematode database and displays fewer missed and duplicated genes than the previous assembly (PX356), but for the most part the BUSCO scores are very similar (see Supplementary Figure S2).

We used full-length transcripts, RNA-seq data from previous *C. remanei* studies, known *Caenorhabditis* proteins, and *ab initio* predictions to annotate the *C. remanei* genome (see Methods). The final annotation contains 26,308 protein-coding genes, which is close to the number of annotated genes in other *C. remanei* strains (Table 1). Each of the genes predicted by AUGUSTUS has been validated by at least one type of evidence: 25,380 genes have hits with known *Caenorhabditis* proteins (23,840 with *C. elegans, C. briggsae, C. latens*), including 25,373 that have matches with the previously annotated genes of *C. remanei;* 19,285 contain known protein domain or functional annotation; 18,662 were supported by RNA-seq data; 8,870 have full-transcript evidence derived from 19,410 high-quality isoforms from the Iso-seq data. In addition, 27 genes were predicted from the full-transcript data.

### Synteny of *C. remanei* and *C. elegans*

We identified 11,160 one-to-one orthologs of *C. remanei* and *C. elegans* protein coding genes, which, after additional filtering on the global-alignment identity, resulted in 9,247 orthologs pairs. Comparing our new chromosome-level assembly to that of *C. elegans* reveals that *C. remanei* and *C. elegans* genomes are in highly synteny despite having a very large number of within-chromosome rearrangements (only 120 of ortholog pairs are located not on homologous chromosomes). The distribution of orthologs across chromosomes is fairly uniform (chromosome I contains 1511 orthologs; II – 1498; III – 1473; IV – 1472; V – 1703; X – 1470). The central domains of autosomes and most regions of the X chromosomes are more highly conserved than the rest of the genome (Figure 1). Orthologs located on X chromosome have greater global identity than ones located on autosomes (W = 532160, p-value = 0.0128).

**Figure 1.**
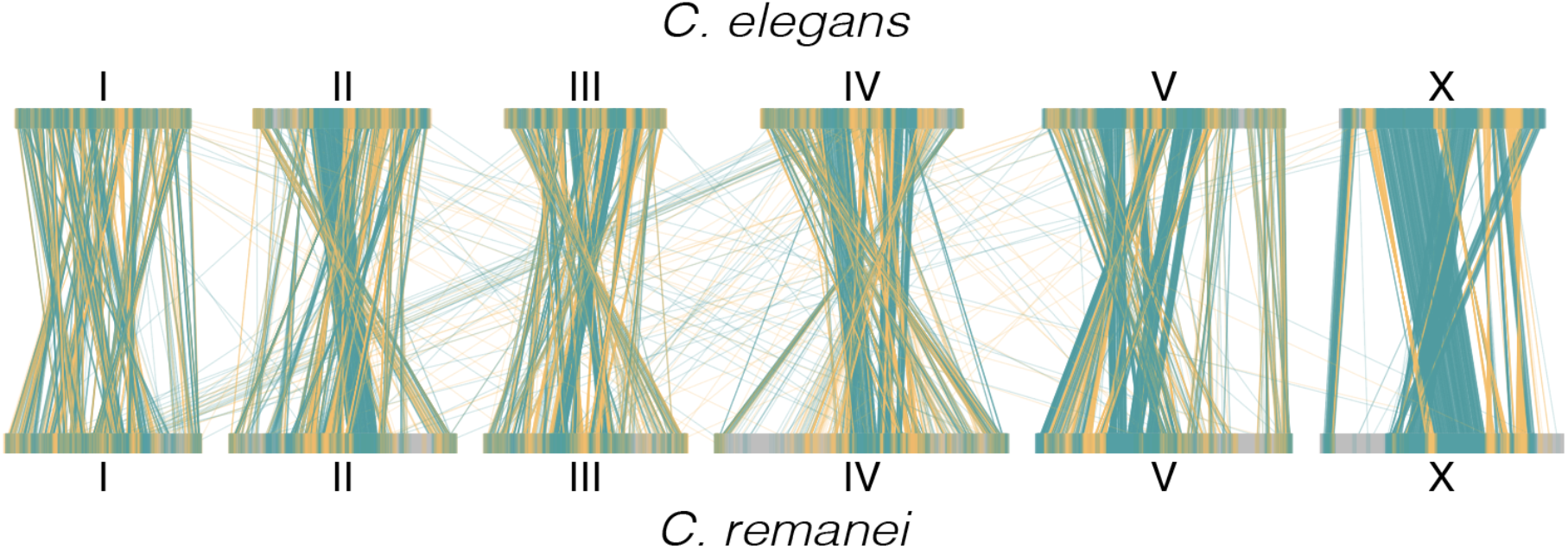
Gene synteny plot. 9,247 one-to-one orthologs of *C. elegans* (the top row) and *C. remanei* (the bottom row). The lines connect locations of the orthologs on the *C. elegans* and *C. remanei* reference genomes. The teal color represents the same orientation of the genes. The orange color shows the inverted orientation.

We chose the orientation of the *C. remanei* chromosomes based on same/inverted direction of nucleotide alignments and 1-to-1 orthologs in *C. elegans.* However, it appears that the ancestral orientation of chromosome III is actually inverted relative to the *C. elegans* standard based on syntenic blocks between *C. briggsae* and *C. remanei* (e.g., *C. elegans* chromosome III has undergone large scale inversion since divergence from the common ancestor of these three species, see the dot-plots on Supplementary figure S3).

### Organization of *C. remanei* and *C. elegans* chromosomes

To compare the genomic organization of *C. remanei* and *C. elegans*, we identified repetitive sequences in the *C. remanei* PX506 genome and the updated reference of *C. elegans* (Yoshimura *et al*. 2019). In total, 22.04% from 124.8 Mb and of the *C. remanei* genome and 20.77% of 102 Mb in the *C. elegans* genome were repetitive. All homologous chromosomes of *C. remanei* are on average 22% longer than corresponding homologous chromosomes in *C. elegans;* the physical size of chromosomes I, II, III, IV, V, and X are 15.3, 15.5, 14.1, 17.7, 21.2, and 18.1 Mb in *C. elegans* and 17.2, 19.9, 17.8, 25.7, 22.5, 21.5 Mb in *C. remanei.* These findings are consistent with the conclusions of (Fierst *et al*. 2015) that the difference in the genome sizes of outcrossing and selfing *Caenorhabditis* species cannot be explained solely by an increase in transposable elements abundance.

In order to identify finer scale patterns displayed across each chromosome, we estimated fractions of exons, introns, repetitive DNA per 100 Kb windows (Figure 2), as well as GC-content, gene counts, and gene fractions (Supplementary Figure S4). In general, *C. elegans* and *C. remanei* display analogous patterns of organization of across all chromosomes. Repetitive DNA was found in greater quantities in the peripheral parts of chromosomes of *C. elegans* (Cohen’s d=1.58, W=232780, p-value < 2.2e-16) and *C. remanei* (Cohen’s d=1.44, W=332810, p-value < 2.2e-16). Repetitive regions of *C. elegans* (VC2010) and *C.remanei* (PX506) genomes are available in Supplementary files S3 and S4.

**Figure 2.**
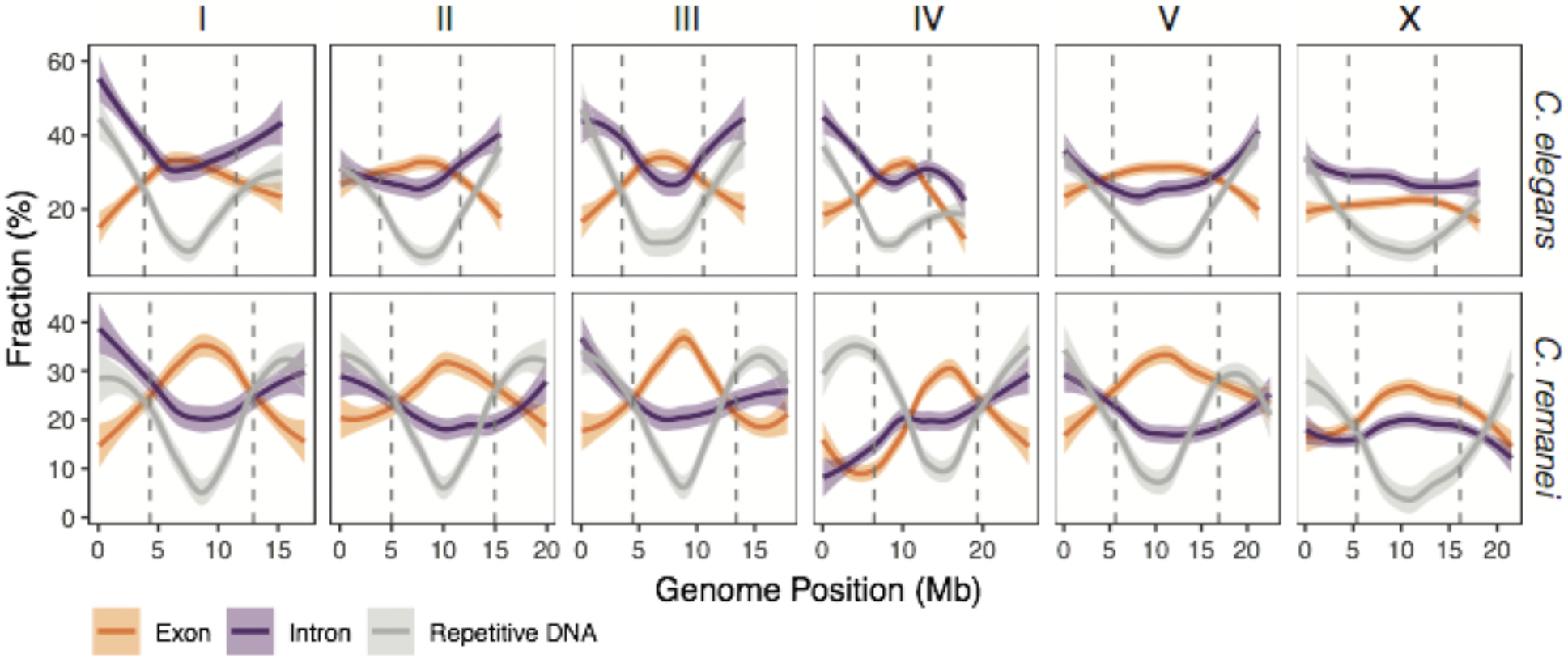
Genomic landscape of genetic elements for *C. elegans* and *C. remanei.* The vertical dashed lines show the boundaries of the “central” domain. Fractions of repetitive DNA, exons, and introns are estimated for 100 Kb windows.

Further, the fractions of the repetitive DNA in both species is negatively correlated with number of genes (r= −0.26 in *C. elegans;* r=-0.45 in *C. remanei)* and the exonic fractions (r= −0.43 and −0.63). There is an inverse positional effect with respect to the number of genes: more genes are located in the central domain than in the peripheral parts of chromosomes in *C. elegans* (Cohen’s d=0.44, W=93950, p-value=2.9e-15) and in the *C. remanei* genome (Cohen’s d=0.72, W=116470, p-value < 2.2e-16), as has been long noted in *C. elegans* (Barnes *et al*. 1995). Both species have a similarly high density of genes (211.2 and 216.2 genes per Mb for *C. elegans* and *C. remanei*, respectively), which is one order of magnitude higher than for humans (Dunham *et al*. 2004). Not surprisingly, then, genes occupy a large fraction of the genome in both *C. elegans* (the mean fraction per 100 Kb is 0.58 with 95% confidence interval 0.569-0.5836) and *C. remanei* (the mean fraction equals 0.44 with 95% confidence interval 0.436-0.452). Genes on the arms have longer total intron size then in the central domains (*C. elegans:* W=53932000, p-value < 2.2e-16; *C. remanei:* W=82764000, p-value < 2.2e-16). GC content, number genes, and gene fraction also differ between central and peripheral parts of chromosomes, as shown in Supplementary figure S4 and Supplementary Table S4.

In both species there is more intronic DNA on the periphery of chromosomes than in their centers (Cohen’s d=0.68, W=177730, p-value < 2.2e-16 for *C. elegans;* Cohen’s d=0.32, W=227220, p-value=1e-06 for *C. remanei*), although for *C. remanei* this pattern is not significant due to the different distributions of introns on chromosomes IV and X (Figure 2). Overall, 28.5% and 27.3% of total intron length consist from repetitive elements in *C. elegans* and *C. remanei.* Additionally, we investigated the transcriptional landscape of *C. elegans* and *C. remanei* genomes on the L1 larval stage, and found that expression of genes in the central domain is very slightly yet significantly larger than gene expression on the peripheral domains (Cohen’s d= 0.06, W=9796800, p-value < 2.2e-16 for *C. elegans;* Cohen’s d=0.04, W=29629000, p-value=2.9e-14 for *C. remanei)*, the chromosome-wise distribution of RPKM is shown on Supplementary Figure S5.

### Similar patterns of within-genome interactions

In examining the pattern of read-mapping of Hi-C data across the *C. remanei* genome, we noted that the central domain of each chromosome appears to be enriched for interactions with the central domains of all other chromosomes (Supplementary Figure S1). To explore this further, we examined the distances of 3D interactions within chromosomes and proportions of inter-chromosomal contacts in *C. remanei* and *C. elegans* genomes. This analysis should be considered to be preliminary as the data is likely noisy since it was obtained from mixed tissues and the *C. remanei* sample was collected from mixed developmental stages (including adult worms), whereas the *C. elegans* results are derived a reanalysis of data from embryos (Brejc *et al*. 2017). At the moment, Hi-C data for different developmental stages of *C. elegans* and/or *C. remanei* is not publicly unavailable.

A total of 12% of the 199.2M of read-pairs mapped to different chromosomes of *C. remanei*, which indicates a high level of potential trans-chromosome interactions. We observed an even higher proportion (32.7% from 123.9 M read pairs) of trans-chromosome contacts in the *C. elegans* sample. When we consider interactions within rather than between chromosomes, we find that the central domains tend to have a larger median distance between interaction pairs compared to the arms. This difference is significant within both species (Figure 3A; Cohen’s d = 1.46, W=39418, p-value < 2.2e-16 for *C. elegans*, and Cohen’s d= 1.74, W=45396, p-value < 2.2e-16 for *C. remanei).*

**Figure 3.**
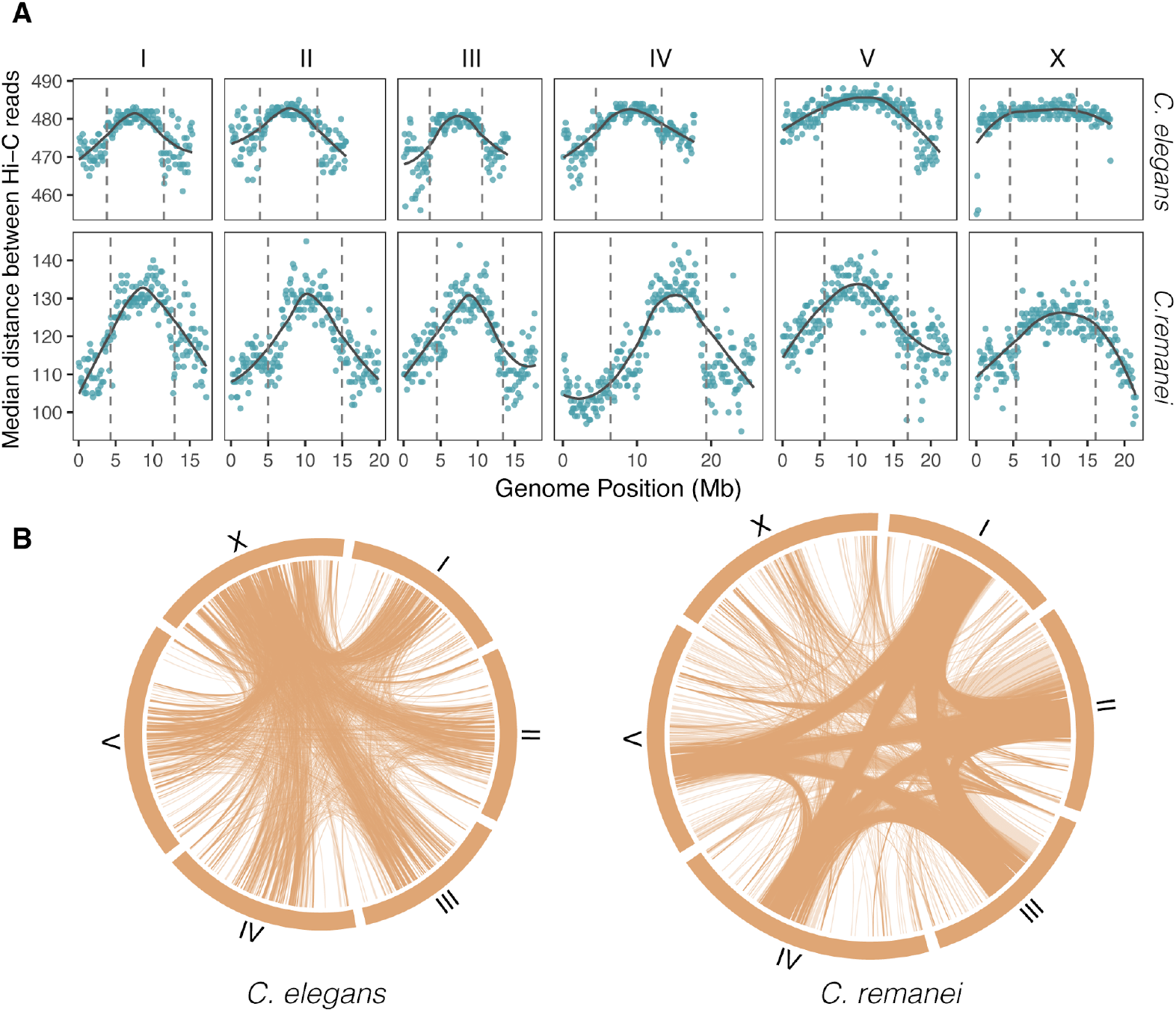
Genome landscape of median distances in Hi-C reads pairs in *C. elegans* and *C. remanei* samples. (A) Distributions of distances between paired reads of cis-chromosome interactions. The vertical dashed lines show the boundaries of the “central” domain. The medians are estimated per 100 Kb windows. (B) Trans-chromosome interactions in *C. elegans* and *C. remanei.* Lines represent contacts between 100 Kb windows. For the *C. elegans* dataset, only contacts with more than 200 pairs of reads are shown; for the C. remanei dataset, only contacts where more than 100 pairs of reads are shown (the *C. remanei* Hi-C dataset is two times smaller than the *C. elegans* dataset). These filters emphasize differences in interaction density/location, the actual total number of interactions is approximately the same for all chromosomes.

Central domains tend to interact with other central domains in *C. remanei* (Figure 3A; 36.2% Center-Center contacts, 40.6% Arm-Center, 19.3% Arms-Arms), but the proportion of Center-Center contacts in *C. elegans* is lower (27.8 % Center-Center, 49.7 % Arm-Center, and 22.5% Arm-Arm). The deviation from the expected uniform distribution of trans-chromosome interactions is larger in *C. remanei* than in *C. elegans* (X-squared = 330220, d.f. = 2, p-value < 2.2e-16 for *C. elegans;* X-squared = 1643800, d.f. = 2, p-value < 2.2e-16 for *C. remanei).* All chromosomes both in *C. elegans* and *C. remanei* sample have almost even number of contacts with other chromosomes (*C. elegans:* I chromosome have 15.1% from all inter-chromosomal contacts, II – 15.3%, III – 14.2%, IV – 17%, V – 19.5%, X – 18.8%; *C. remanei:* I – 16.1%, II – 16.7%, III – 16.1%, IV – 18.2%, V – 18.2%, X – 14.7%). However, if we focus specifically on windows with localized contacts, we see that within *C. remanei* interactions are more dispersed on X and V chromosomes and there are areas of thick contacts in the central parts of autosomes, whereas, in the *C. elegans* sample all chromosomes actively interact (Figure 3B).

## Discussion

We have generated a high-quality reference genome of the *Caenorhabditis remanei* line PX506, which is now one of the five currently available chromosome-level assemblies of *Caenorhabditis* nematodes of the Elegans supergroup, including three selfing species, *C. elegans (C. elegans* Sequencing Consortium 1998), *C. briggsae* (Stein *et al*. 2003), and outcrossing *C. inopinata* (Kanzaki *et al*. 2018) and *C. nigoni* (Yin *et al*. 2018). *C. remanei* is an outcrossing nematode with high genetic diversity in comparison with *C. elegans, C. briggsae* and *C. tropicalis* (Jovelin *et al*. 2003; Cutter *et al*. 2006). Therefore, to reduce the diversity and improve the quality of assembly, we constructed a highly-inbred line from wild isolates collected from a forest near Toronto (see Methods). As expected, we assemble a genome consisting of six chromosomes, each of which is largely syntenic at a macro level with the genome assemblies from the other *Caenorhabditis* species. The difference in the genome lengths between *C. elegans* and *C. remanei* is quite large, from 102 Mb to well over 124 Mb. However, this degree of size variation appears to be typical for *Caenorhabditis* nematodes. For example, Stevens *et al*. (2019) showed that the genome sizes across the genus can vary from 65 Mb to 140 Mb and that, overall, the size of the genome correlated with the number of genes but not necessarily the mode of reproduction.

This is the third *C. remanei* genome assembly generated by our group (PX439, PX356, and PX506; Table 1). The two previous chromosome-scale assemblies of other *C. remanei* strains (PX439, PX356) were constricted with Illumina data and multiple matepair libraries (Fierst *et al*. 2015). However, the *C. remanei* genome has extended repetitive regions that failed to assemble using short reads. Further, strong segregation distortion among strains made it very difficult to construct the genetic map and definitively align shorter contigs to specific putative. In this study, we used deep PacBio sequencing and Hi-C linkage information to overcome the repetitive regions and achieve better assembly characteristics. The combination of long-read and linkage data is a powerful toolset to produce chromosome-level assemblies, which is currently being increasingly used in a large number of species (e.g., Gordon *et al*. 2016; Gong *et al*. 2018; VanBuren *et al*. 2018; Low *et al*. 2019).

In addition to genome assembly, we performed annotation of the new *C. remanei* reference genome, using full-length transcripts data (Iso-Seq), which has proven to be an effective technique to create a high-quality annotation (Gonzalez-Garay 2016), shortread transcriptome sequencing, protein sequences of related species, and a hybrid annotation pipeline. To validate predicted gene models, we additionally used the previous annotation of *C. remanei* since it was manually curated and mostly was supported by RNA-seq data (Fierst *et al*. 2015). The genes that were not present in the previous annotation are supported by other lines of evidence, including genes predicted from the Iso-Seq data. We find a total of 26,308 genes in the *C. remanei* genome, a slight increase over previous estimates (Fierst *et al*. 2015) and a reconfirmation that *C. remanei* appears to have more genes than the selfing species.

We compared the genomic organization of *C. remanei* and *C. elegans* using the last available version of the VC2010 *C. elegans* genome, which is based on a modern strain derived from the classical N2 strain and which led to an enlargement of the N2-based genome by an additional 1.8 Mb of repetitive sequences (Yoshimura *et al*. 2019). *C. remanei* and *C. elegans* genomes are in highly synteny in spite of multiple intrachromosomal rearrangements (Figure 1). We observed many more intra-than interchromosomal rearrangements, which is consistent with first comparative observations of the *C. elegans* and *C. briggsae* genomes, which saw a 10-fold difference in these rates (Stein *et al*. 2003). This overall pattern remains consistent even when comparing *C. elegans* to more distantly related genera of nematodes (Guiliano *et al*. 2002; Whitton *et al*. 2004; Mitreva *et al*. 2005).

One plausible explanation for this pattern is that the low rate of interchromosomal translocations is generated by the multilevel control of meiotic recombination in *Caenorhabditis.* Pairing of chromosomes during meiosis in *C. elegans* is started from specific regions (“pairing centers”) located on the ends of homologous chromosomes (MacQueen *et al*. 2005; Tsai and McKee 2011), followed by chromosome synapsis via assembly of synaptonemal complex along coupled chromosomes (MacQueen *et al*. 2002; Rog and Dernburg 2013). Crossovers in *C. elegans* can be formed only between properly synapsed regions (Lui and Colaiácovo 2013; Cahoon *et al*. 2019). Taken together, these molecular mechanisms permit meiotic recombination only between homologues region linked in *cis* to the “pairing centers,” which presumably reduces the number of interchromosomal rearrangements, thereby resulting in the evolutionary stability of the nematode karyotype (Rog and Dernburg 2013).

The central domains of autosomes and a large portion of the X chromosome have more extended conservative regions between *C. remanei* and *C. elegans*. The similar pattern has been observed in comparative genomic studies of *C. elegans* and *C. briggsae* (Stein *et al*. 2003; Hillier *et al*. 2007). Apparently, the stability and conservation of the central regions is also connected to the recombinational landscape, as the central half chromosomes in *C. elegans* display a recombination rate that is several times lower than that observed on chromosome ends (Rockman and Kruglyak 2009), *C. briggsae* (Ross *et al*. 2011), as well as in *C. remanei* (Teterina et al., in prep), without definitive hotspots of recombination (Kaur and Rockman 2014). The rate of recombination on the X chromosome is less than that on autosomes (Bernstein and Rockman 2016), and because of the XX/X0 sex determination system of nematodes, the population size of the sex chromosome is ¾ that of the autosomes (Wright 1931). So orthologs of *C. elegans* and *C. remanei* located on X chromosome are more conserved on average, likely because selection against deleterious mutations on the sex chromosome is greater than on autosomes (Montgomery *et al*. 1987; Coghlan and Wolfe 2002).

The chromosomes of *C. elegans* (Consortium 1998) and *C. remanei* also have a very similar pattern of gene organization, with a central region (the “central domain” or “central gene cluster;” (Barnes *et al*. 1995) characterized by high gene density, shorter genes and introns, lower GC-content (Supplementary Figure S4 and Table S4) and almost two times lower abundance of repetitive elements compared to chromosome “arms”. Repetitive elements in *C. elegans* and *C. remanei* are more abundant in the peripheral parts of chromosomes and, respectively, leaving a reduced fraction of protein coding genes in those regions. About 28% of the total intron length in these nematodes are occupied by TEs, which could partially explain the increase of the gene lengths and intron fractions on the “arms”. The positive correlation of intron size with recombination rate and transposable elements has been previously observed in *C. elegans* (Prachumwat *et al*. 2004; Li *et al*. 2009b). The “central gene cluster” and transposable elements enriched in the “arms” is common and is likely the ancestral pattern observed in *C. elegans, C. briggsae, C. tropicalis* and *C. remanei*, yet distinct in *C. inopinata* (Woodruff and Teterina 2019).

Use of Hi-C data in the genome assembly allows us to perform a preliminary analysis of the three-dimensional chromatin organization across mixed developmental stages in *C. elegans* and *C. remanei.* The central domains show more cis-chromosome interactions than the peripheral parts of chromosomes in *C. remanei* (Figure 3). In *C. elegans* variation in interaction intensity across the chromosome is somewhat less perceptible, probably because of minor differences in the fractions of genes on the central domains versus arms. In both species, central regions show more distant interactions then arms. All chromosomes have numerous trans-chromosome interactions that are more tightly localized in the central regions. This pattern can be explained both by density of genes in the central domains but also by technical issues with mapping of the reads to the repetitive regions. In contrast to the autosomes, the pattern of trans-chromosome activity is more dissimilar in *C. elegans* and *C. remanei*. This could be caused by species-specific differences or by the fact that the developmental stages of the samples do not strictly correspond for the two species (the *C. elegans* dataset used early embryos whereas the *C. remanei*sample included all stages of the life cycle). In this case, both X chromosomes are active in hermaphrodites (XX) but their activity is reduced in half by dosage compensation mechanism in all tissues in *C. elegans* after the 30-cell stage (gastrulation) (Meyer 2005; Strome *et al*. 2014; Crane *et al*. 2015; Brejc *et al*. 2017). The presence of individuals at the early developmental stages could therefore potentially affect the extent of interactions with X chromosome observed within the *C. elegans* sample. Dosage compensation suppresses gene expression on both X chromosomes, modulates chromatin conformation by forming topologically-associated domains, and partially compresses both X chromosomes (Meyer 2010; Lau *et al*. 2014; Brejc *et al*. 2017). All of these structural changes could potentially affect relative intensity and availability of interactions between the X chromosome and autosomes.

What might drive these inter-chromosomal interactions? Cis-and trans-chromosome interactions could mediate transcriptional activity through co-localization of transcriptional factors on gene regulatory regions (Miele and Dekker 2008; Pai and Engelke 2010; Maass *et al*. 2019). Genome activity and spatial organization of a genome is a dynamic property, and chromatin accessibility in *C. elegans* is tissue-specific, change over developmental time (Daugherty *et al*. 2017; JӒnes *et al*. 2018). However, *C. elegans* tends to have active euchromatin in the central parts of chromosomes and silent heterochromatin in the arms, which are anchored to the nuclear membrane (Ikegami *et al*. 2010; Liu *et al*. 2011; Mattout *et al*. 2015; Solovei *et al*. 2016; Cabianca *et al*. 2019). This pattern of regulation is consistent with the pattern of interactions that we observe. Nevertheless, much more work needs to be conducted, particularly aimed at stage and tissue specific effects, before the role and dynamics of spatial chromosome interaction in *Caenorhabditis* can be fully revealed.

Overall, then, despite numerous within-chromosomal rearrangements, *C. elegans* and *C. remanei* show similar patterns of chromosomal structure and activity. The chromosome-level assembly of *C. remanei* presented here provides solid new platform for experimental evolution, comparative and population genomics, and the study of genome function and architecture.

## Acknowledgments

We thank members of the Phillips lab for helpful discussions and the University of Oregon Genomics and Cell Characterization Core Facility (GC3F) for advice and support. We are grateful to Tim Ahearne and Scott Sholtz for generating the *C. remanei* PX506 inbred line. This work was supported grants from the National Institutes of Health (R01GM102511, R35GM131838) to P.C.P.

## Supplementary Material

**Supplementary Figure S1**. Hi-C Link density histogram of the *C. remanei* PX506 inbred line generated by Dovetail Genomics. https://doi.org/10.6084/m9.figshare.11466966.v1

**Supplementary Figure S2**. BUSCO-analysis of assembly completeness for *C. briggsae, C. elegans, C. remanei* genomes with (A) Metazoa database od9 and (B) Nematoda database od9. https://doi.org/10.6084/m9.figshare.11466975.v1

**Supplementary Figure S3**. Synteny of (A) *C. elegans* and *C. remanei*, (B) *C. briggsae* and *C. remanei.* https://doi.org/10.6084/m9.figshare.11466987.v1

**Supplementary Figure S4**. Genomic landscapes of *C. elegans* and *C. remanei* (A) GC-content, (B) gene fractions, and (C) the number of genes. All values are estimated per 100 Kb windows. https://doi.org/10.6084/m9.figshare.11467002.v1

**Supplementary Figure S5**. Genomic transcriptional landscape on L1 larval stage in *C. elegans* (top) and *C. remanei* (bottom). The heatmap is based of RPKM values per gene. https://doi.org/10.6084/m9.figshare.11470389.v1

**Supplementary Tables S1, S2**, and **S3** with additional assembly and PacBio statistics for the *C. remanei* genome assembly generated by Dovetail Genomics are included in the file **Supplementary_File1.docx**. **Supplementary Table S1**. FALCON and HiRise assembly statistics. **Supplementary Table S2**. Summary statistics for the PacBio data used in the study. **Supplementary Table S3**. Additional statistics for the HiRise assembly. https://doi.org/10.6084/m9.figshare.11466963.v1

**Supplementary Table S4**. Comparative statistics of “Arms” and “Centers” domains of *C. elegans* and *C. remanei* chromosomes is in **Supplementary_File_S2.docx**. https://doi.org/10.6084/m9.figshare.11466951.v1

**Supplementary File S3.** Repetitive regions of the *C. remanei* genome (PX506) in the bed file format. https://doi.org/10.6084/m9.figshare.11470830.v1

**Supplementary File S4.** Repetitive regions of the *C. elegans* genome (VC2010) in the bed file format. https://doi.org/10.6084/m9.figshare.11470833.v1

